# The emergence of network structure, complementarity and convergence from basic ecological and genetic processes

**DOI:** 10.1101/007393

**Authors:** Francisco Encinas-Viso, Carlos J. Melián, Rampal S. Etienne

## Abstract

Plant-animal mutualistic networks are highly diverse and structured. This has been explained by coevolution through niche based processes. However, this explanation is only warranted if neutral processes (e.g. limited dispersal, genetic and ecological drift) cannot explain these patterns. Here we present a spatially explicit model based on explicit genetics and quantitative traits to study the connection between genome evolution, speciation and plant-animal network demography. We consider simple processes for the speciation dynamics of plant-animal mutualisms: ecological (dispersal, demography) and genetic processes (mutation, recombination, drift) and morphological constraints (matching of quantitative trait) for species interactions, particularly mating. We find the evolution of trait convergence and complementarity and topological features observed in real plant-animal mutualistic webs (i.e. nestedness and centrality). Furthermore, the morphological constraint for plant reproduction generates higher centrality among plant individuals (and species) than in animals, consistent with observations. We argue that simple processes are able to reproduce some well known ecological and evolutionary patterns of plant-animal mutualistic webs.

## Introduction

Since Darwin's book “On The Origin of Species” (Darwin, 1862*b*), the idea of coevolution ^1^ has sparked interest from biologists trying to understand how species interactions generate trait changes. The first clear indication of coevolution was Darwin's moth example (Darwin, 1862*a*) showing that the long corolla from the orchid *Angraecum sesquispedale* could only be reached by a pollinator species with a similar proboscis length. However, much later Janzen (1980) argued that this amazingly high specialization between plants and animals was not the only example of coevolution. He explained that coevolution can also be the product of multiple-species interactions, a term that he coined “diffuse coevolution”. Diffuse coevolution means that selection on traits is determined by the interaction of all species in the community and not only based on pair-wise interactions. This is based on the idea of pollination or dispersal “syndromes”, where plants have a set of traits that attract a specific group of pollinator or animal seed-disperser species.

Later, the idea of “diffuse coevolution” was related to patterns of nestedness detected across biogeographic regions in mutualistic networks. Nestedness, defined as a non-random pattern of interactions where specialist species interact with proper subsets of more generalist species puts the concept of “diffuse coevolution” in a more quantitative context (Bascompte et al., 2003). Nestedness patterns have been shown to provide information about the underlying network dynamics. For example, nestedness is associated with stability and coexistence of species in a community (Bastolla et al., 2009; Okuyama and Holland, 2008).

Several studies have modelled coevolutionary dynamics in mutualistic systems of a few species (Ferriere et al., 2007; Law et al., 2001; Ferdy et al., 2002; Gomulkiewicz et al., 2003; Jones et al., 2009), particularly highly specialized (i.e. obligatory mutualists) systems of plant-animal interactions, such as the fig-fig wasp mutualism (Bronstein et al., 2006). These studies have determined the ecological conditions for coevolutionary stable systems (i.e. coESS) (Law et al., 2001; Jones et al., 2009). However, more complex cases of evolution involving multispecific interactions in the context of quantitative genetics and explicit speciation mechanisms remain unexplored.

There are two trait-based patterns in plant-animal mutualistic networks that provide evidence for niche-driven and coevolutionary processes shaping these webs: evolutionary *complementarity* and *convergence* (Bascompte and Jordano, 2007). Complementarity describes that there is selection for trait matching between plant and animal traits (e.g. corolla length-proboscis length, frugivore body mass-seed size) (Rezende et al., 2007; Bascompte and Jordano, 2007). Therefore, complementarity seems the clearest explanation of reciprocal evolution (i.e. coevolution). Convergence is consistent with observed trait similarity among evolutionarily distantly related species of the same guild (e.g. pollinators with similar proboscis length) and is assumed to be caused by selective pressures and developmental constraints (Bascompte and Jordano, 2007). Evolutionary convergence in plant-animal mutualisms partly explains the formation of ‘syndromes’ produced by the presence of specific mutualist partner species (Bascompte and Jordano, 2007; Howe and Smallwood, 1982; Waser et al., 1996). For example, plant species with a specific corolla morphology may determine the evolutionary convergence of pollinator species traits (Jousselin et al., 2003). Guimaraes et al. (2011) studied a coevolutionary model of mutualistic webs where selective pressures came only from mutualistic partners and found that coevolution promotes complementarity and convergence supporting the idea that selection through niche-driven mechanisms (i.e. the biotic environment) is mainly responsible for the observed patterns. However, non-selective causes can also produce evolutionary convergence (Losos, 2011).

Krishna et al. (2008) and Canard et al. (2012) have shown that random fluctuation of species abundance (i.e., ecological drift) can explain some of the topological properties in mutualistic and trophic webs, respectively. These studies do not take into account explicitly the genetics of quantitative traits and speciation dynamics. The question then arises whether models that describe quantitative trait dynamics with explicit genetics and speciation in the context of random fluctuations of species can generate simultaneously the evolution of convergence, complementarity and network topology observed in real plant-pollinator webs.

Recently, various neutral eco-evolutionary models have started to consider genetics explicitly and more realistic assumptions about the speciation process (de Aguiar et al., 2009; Melián et al., 2012). These models, which consider intraspecific variation and explicitly incorporate three of the main evolutionary forces (mutation, recombination and drift (Lynch, 2007; Vellend, 2010)), can predict biodiversity patterns well. Furthermore, these models use a common theoretical framework based on the neutral theories of evolution (Kimura, 1983) and ecology (Hubbell, 2001). They allow testing model predictions with available data on diversity, species traits, spatial distribution and genetics. The progress in this area is rapid, but it is still in its early stages.

Here, we develop an individual-based stochastic model of plant-pollinator interactions that considers explicit genetics, phenotype expression and spatial structure of sexually reproducing individuals, to study the eco-evolutionary dynamics of plant-pollinator webs. We find emergence of plant-pollinator network topological properties such as nestedness and centrality, and the evolution of trait convergence and complementarity. We argue that basic ecological and genetic processes in combination with physical constraints of plant-pollinator interactions, can generate observed plant-pollinator network topology and the evolutionary patterns of plant-pollinator traits.

## The model

We consider the eco-evolutionary dynamics of plants (*P*) and animal pollinators (*A*). These two guilds interact mutualistically: plants need the presence of pollinators and vice versa to reproduce. Hence the mutualism is obligatory for both partners.

### General eco-evolutionary dynamics

Our model is a stochastic individual-based model with overlapping generations and zero-sum birth-death dynamics. The population consists of *J_P_* and *J_A_* haploid gonochoric (i.e. separated sexes) individuals for plants and animals, respectively; with explicit binary genomes of size *L.* Each plant and animal population reproduces sexually and is spatially structured. The reproduction of each guild is done in turns (i.e. asynchronically). The individual-based events occur in the following order: an individual is randomly selected to die and then a female individual is randomly chosen among all females within a distance *d_max_*of the dead individual's position to mate. Thus, death and reproduction events only occur at a local scale to reflect limited dispersal.

There are two conditions for sexual reproduction: 1) the geographic distance *d_ij_* between two individuals (plant or animal), a female *i* and a male *j*, from the geographic distance matrix *D* has to be lower than the maximum geographic distance *d_max_* (*d_ij_* < *d_max_*). In case there are no potential mates, a different female is randomly chosen until a potential mate is found. We have two geographic distance matrices: *D^P^* and *D^A^* for plants and animals, respectively. 2) the genetic similarity *q_ij_* between two individuals (defined below) has to be higher than the minimum genetic distance *q_min_* to be able to mate and leave viable offspring (hence individuals mate assortatively). The genome of each individual is represented by a sequence of *L* loci, where each locus can be in two allelic states, +1 or −1. Each individual *i* in a population of size *J* is represented as a vector: 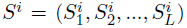, where 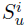 is the *u^th^* locus in the genome of individual *i*. The genetic similarity between individuals is calculated as the sum of identical loci across the genome:

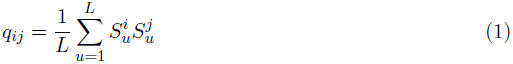

where *q_ij_* ∈ {−1, 1}. The offspring born from this mating is dispersed within the geographic distance, *d_max_*, and will occupy the geographic position of the just deceased individual.

The genome of the offspring is obtained by a block cross-over recombination of the female genome *S^i^* and male genome *S^j^*, where a locus *l* in the genome of the parents is randomly chosen partitioning the genome of each individual in two blocks. All genes beyond that locus *l* in either organism genome are swapped between the two parents and eventually form two new genomes. One of the two new genomes is randomly chosen from a uniform distribution for the offspring. The offspring's genome undergoes mutations at mutation rate *µ*. Figure 1 describes the model, including the recombination-mutation process.

**Figure 1:**
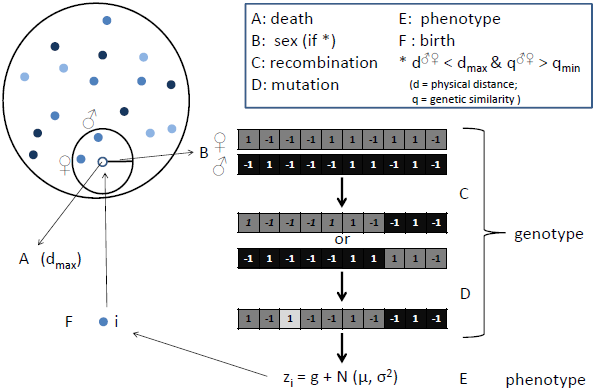
General description of the model. Each time step in this model is completed after a death-birth cycle (from A to F). Individuals are represented as filled circles scattered across space and the variation of blue colors represents their variation of phenotypes. The model is divided into different events at each time step: (A) an individual *k* is randomly selected to die and leaves an empty location in the landscape. (B) a female individual *i* is randomly selected if *d_kf_* < *d_max_* and this female *f* will mate and reproduce with a male individual *j* if conditions of mating are met (*d_ij_* < *d_max_* and *q_f j_* > *q_min_*) (additional mating conditions depend on the guild, but always required the presence of a mutualistic partner, see Methods section). (C) The recombination process. Genomes are composed of *L* loci where each locus can be in two allelic states (−1, 1) and undergo block cross-over recombination of the female genome (dark gray) and male genome (black), where a position *l* in the genome of the parents is randomly chosen partitioning the genome of each individual in two blocks. In this example the genome it is split into parts of equal length. All genes beyond the *l* locus in either organism's genome is swapped between the two parents and two new genomes are formed. (D) One of the two new genomes is randomly chosen for the offspring and it might undergo mutation (light gray). (E) The phenotype expression of newborn individual *i* is *z_i_* = *g_i_* + *∈*. (F) The newborn *i* will occupy the site of the dead individual *k* within the area *d_max_*.

At the beginning of the simulations all individuals are genetically identical (*q* = 1), hence they are all able to mate with one another. We can visualize the genetic similarity between individuals of a guild as an evolutionary spatial graph (Melián et al., 2010), where nodes correspond to individuals and the length of edges correspond to the geographic distance between a pair of genetically similar (*q_ij_* > *q_min_*) individuals. At the beginning of the simulation this leads to a fully connected graph under an evolutionary process with mutation, recombination and dispersal. The connectance of the graph will decrease when species are formed (i.e. speciation). Here, we define a species as a group of genetically related individuals, where two individuals in sexual populations can be conspecific while also being incompatible, as long as they can exchange genes indirectly through other conspecifics. This is the definition of ‘ring species’ (de Aguiar et al., 2009; Melián et al., 2010).

The speciation process in this model is similar to previous neutral speciation models with explicit genetics (de Aguiar et al., 2009; Melián et al., 2010). Individuals become more and more genetically divergent through the mutation and recombination process and the spatial segregation. This will finally produce the formation of two genetically incompatible clusters of individuals, i.e. two species. This speciation process, also called ‘fission-induced’ speciation (Melián et al., 2012), goes on with the formation of more clusters and genetic divergence between individuals of different species. However, the diversification dynamics will fluctuate due to random extinctions (death of last individual of a species). A stochastic balance between speciation and extinction is eventually reached giving the final steady-state of the metacommunity (Melián et al., 2012).

### Quantitative traits

The quantitative trait (*z*) of each individual is determined by additive genetic effects of the genome (*g*) (i.e. no epistasis) plus a normally distributed environmental effect 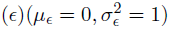. Thus, *z_i_* = *g_i_* + *∈* determines the phenotype or quantitative trait (*z_i_*) of each individual. The genetic component (*g_i_*) of an individual *i* is:

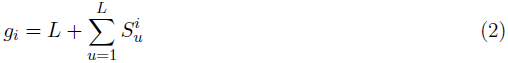

calculated as the sum of alleles across the genome (Kondrashov and Shpak, 1998) plus the number of loci to avoid negative trait values. If we sample genomes of size *L* from a uniform distribution, the distribution of genetic values would have mean *L* and a variance given by the algebraic sum of allelic values. We assume two quantitative traits, one for each guild: proboscis length 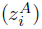 in pollinators and corolla length 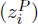 in plants.

### Phenotypic similarity

We measured phenotypic similarity between individuals of the same guild and of different guilds to study the relationship between genotypic and phenotypic similarity. The phenotypic similarity (*p_ij_*) between an inidividual *i* and an individual *j* is defined as:

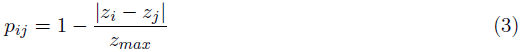

where *z_i_* and *z_j_* are the phenotypic values of individuals *i* and *j*, respectively; and *z_max_* is the maximum value of the phenotype distribution *Z* of the whole metacommunity. Thus, each pair-wise comparison, *p_ij_* ∈ {0, 1}, is an element of the phenotypic similarity matrix *P*.

### Evolutionary convergence and complementarity

We define evolutionary convergence as the similarity between average species phenotypes from distantly related species. We assume that two species are distantly related, in phylogenetic terms, if they do not come from a direct common ancestor, i.e. they are not sister species. To exclude sister species from the analysis we need to calculate the average genetic similarity among species of the same guild. The average genetic similarity between a species *k* and a species *l* is:

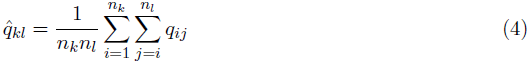

where *q_ij_* is the genetic similarity between an individual *i* of species *k* and an individual *j* of species *l*, and *n_k_* and *n_l_* are the absolute abundances of species *k* and *l*, respectively. The elements *q̂_kl_* will form the matrix *Q_s_* ∈ {*q̂*_*kl*_} from which the sister species of each species in the guild can be identified.

To calculate evolutionary convergence we need to know the average phenotypic similarity between two species. We define phenotypic similarity between species *k* and *l* as:

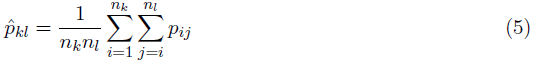

which is analogous to the definition of eq. 4. This will build a species phenotypic matrix *P_s_* ∈ {*p̂*_*kl*_}.

We then focus on each species in turn and exclude its sister species to avoid cases of parallel evolution to calculate the number of convergences related to the focal species. We define a focal species *k* and a non-sister *l* species to be convergent if phenotypic similarity between them is higher than between focal and sister species (*p̂_k,sister_* < *p̂_kl_*) and higher than 0.95 (*p̂_kl_* > 0.95). A simple example to understand the calculation of convergences is illustrated in Figure 2. With only three species, only one convergence is possible after excluding the sister species. Naturally, the number of convergences potentially increases with the number of species present. For example, if we have ten species and we exclude one of them as sister species, we have nine species to calculate convergence with. If we find that two out of nine species are phenotypicallly similar enough to the focal species, we count two (out of nine, ∼ 22%) convergences. Thus, contrary to Guimaraes et al. (2011) we use both genetic divergence and phylogenetic relatedness for the estimation of evolutionary convergences, in order to avoid cases of parallel evolution^2^ (Losos, 2011).

**Figure 2:**
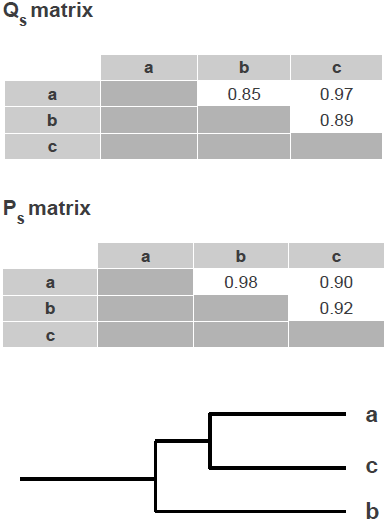
Evolutionary convergence calculations. Convergence is calculated with the species genetic similarity matrix *Q_S_* ∈ {*q̂_kl_*} and the species phenotypic similarity matrix *P_S_* ∈ {*p̂*_*kh*_}. This figure illustrates a simple example of evolutionary convergence where there are only three species in a guild (*a*, *b* and *c*). The upper matrix (*Q_S_*) shows species *a* and *c* are genetically closely related, *q̂_ac_* = 0.97, while genetically distant from species *b* (*q̂_ab_* = 0.85, *q̂_cb_* = 0.89). A clear description of these genetic relationships can be represented with a cluster tree or dendrogram, as shown in the lower part of the figure. Thus, we establish that species *a* and *c* are sister species. The species phenotypic similarity matrix *P_S_* shows that species *a* and *b* are phenotypically highly similar (*p̂_ab_* = 0.98) and highly genetically dissimilar (*q̂_ab_* = 0.85) (i.e. more than the average intraspecific genetic similarity or sister species 0.97), indicating an event of evolutionary convergence.

Evolutionary complementarity is easier to calculate because it does not involve the calculation of the genetic similarity matrix. We only need to estimate the phenotypic similarity between plant and animal species. We do this in the same way as for evolutionary convergence: we calculate the phenotypic similarity matrix *P_PA_* ∈ {*p̂_kh_*} between plant and animal average species traits and the condition for complementarity is that the similarity between a plant species *k* and an animal species *h* should be *p̂_kh_* > 0.95. To visualize the genetic relatedness between species we constructed clustering trees using Euclidean distance with the Python library ETE 2.01 (Huerta-Cepas et al., 2010).

### Plant-animal interactions

In addition to the genetic and geographic constraints for mating, we consider other mating conditions that are different for each guild. These conditions describe the mutualistic interaction between plants and animals and their spatial constraints for interaction. We therefore specify another geographic distance matrix *D^PA^* to describe the geographic distance between plant and animal individuals. Plant-animal mutualistic interactions are here described as follows: plants benefit from the presence of specific pollinators that are able to pollinate them and animals benefit from the presence of plants that provide resources for them. Thus, we have two extra conditions for mating: 1) Plants need the presence of animal pollinators within a close distance 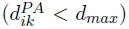 and with a larger or equally-sized proboscis than the corolla of a plant: *z_c_* ≤ *z_p_*. This corresponds to a physical or morphological constraint for individual interactions observed between plant and pollinator species (Stang et al., 2009, 2006). 2) Animals need the presence of plants within a close geographic distance 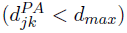. The conditions are illustrated in Figure 1.

Our model allows bookkeeping of who is interacting with whom, i.e. this means we can record exactly which plant and animal individuals are interacting. This bookkeeping not allows comparison with high-resolution data of interactions, as in some plant-pollinator studies (Gómez et al., 2011; Gómez and Perfectti, 2012), but, more importantly for our current aim, enables us to identify the different constraints on the evolution and final topology of the network. We record the identity of the mutualistic partners during the reproduction process for plants and animals after reaching the steady-state to reconstruct the plant-animal interaction network.

### Network topology

We measured three topological properties of plant-animal mutualistic networks: nestedness, connectance and centrality. The topological measurements were applied to the networks at the final steady-state of the simulation.

#### Nestedness

Nestedness describes a non-random pattern of species interactions where specialist species interact with proper subsets of more generalist species (Bascompte et al., 2003). We estimated nestedness using the NODF algorithm developed by (Almeida-Neto et al., 2008) because of its statistical robustness. NODF is based on standardized differences in row and column fills and paired matching of occurrences.

#### Connectance

Connectance measures the proportion of realized interactions (i.e. links) among all possible interactions in a network and is defined as 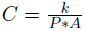, where *k* represents the number of realized interactions between plant and animal species and *P* and *A* represent the number of plant and animal species, respectively, in the network (Jordano et al., 2003).

### Centrality and node redundancy

To explore the topology of the network at the individual level we calculated three different centrality metrics: *degree centrality* (DC), *closeness centrality* (CC) and *betweenness centrality* (BC). These topological metrics are commonly used in social network analysis to study the importance of some nodes in the network, e.g. for the flow of information (Borgatti, 2005) or spread of diseases (Klovdahl, 1985). In ecological networks they have been used to describe the topology of individual-based interaction networks (Gómez and Perfectti, 2012; Gómez et al., 2011) and to identify keystone species that maintain the cohesiveness of the network (Jordán et al., 2007; González et al., 2010).

Degree centrality (DC) is defined as the fraction of nodes connected to a specific node (Borgatti and Halgin, In press). This metric provides a description of how well connected the individuals are. Closeness centrality (CC) measures the distance of a node to all the other nodes in the network; a node with high CC can potentially interact with any other node in the network (Borgatti and Halgin, In press; Newman, 2003). Betweenness centrality is the number of shortest paths between two nodes that pass through a specific node (Borgatti and Halgin, In press; Goh et al., 2003). Therefore, individuals (nodes) with high BC act as bridges, connecting one part of a network to another, maintaining the cohesiveness of the network. We also measured the average clustering coefficient (ACC) and node redundancy (NR) as complementary metrics of centrality. The average clustering coefficient computes the average probability, for any given node chosen at random, that two neighbors of this node are linked together (Latapy et al., 2008), hence it provides evidence of modularity in the network (Olesen et al., 2007). Node redundancy is defined as the fraction of pairs of neighbors of a specific node that are both linked to other nodes (*N R* ∈ {0, 1}). Therefore, higher levels of node redundancy indicate that most nodes in the network share similar partner individuals and hence the elimination of those nodes from the network will not greatly affect the topology. All centrality and node redundancy computations were performed using the Python library NetworkX (Hagberg et al., 2008).

### Simulations

We simulated a population size of *J* = 10^3^ individuals for each guild and a genome size *L* = 150 loci. Larger population sizes (10^4^, 10^5^) are possible, but they are constrained by computational time. Initially all animal individuals have a higher phenotypic trait value than plant individuals (*Z_c_* ≤ *Z_p_*) to assure that plant mating conditions are met at the beginning of the simulation. Geographic distance between each pair of individuals *i* and *j*, *d_ij_*, was calculated as follows: 1) Euclidean coordinates of a two-dimensional space (*x_i_*, *y_i_*) were sampled from a uniform distribution (*x_i_* = [0, 1]*, y_i_* = [0, 1]) for each individual in the metacommunity. 2) Using these coordinates (*x_i_*, *y_i_*) we calculated a matrix of relative Euclidean distances between the individuals (*d_ij_*). This procedure is repeated for each of the geographic distance matrices (*D_PA_*, *D_PP_*, *D_AA_*). The simulation lasted for 2×10^3^ generations, where a generation is an update of *J* time steps. Steady-state was verified by checking the constancy of speciation events during the last 100 generations. We explored a range of parameter combinations with mutation rate, *µ* ∈ {10^−4^, 10^−2^}, minimum genetic similarity *q_min_* = 0.97 and maximal distance *d_max_* = 0.3. We implemented the model in Python (and tested in IPython (Pérez and Granger, 2007)) and graphics were produced using the Python library Matplotlib (Hunter, 2007).

## Results

Mutation rates have an strong effect on the diversity dynamics by affecting the two types of speciation in this model: “mutation-induced” and “fission-induced” speciation. For large mutation rate (*µ* > 10^−2^), speciation is predominantly mutation-induced, resulting in the formation of a species consisting of a single individual. Most of these mutation-induced species are likely to go extinct because of their low initial abundance. For lower mutation rates (*µ* ∈ {10^−4^, 5 × 10^−3^}) speciation was predominantly “fission-induced”, i.e. by slow genetic divergence between individuals, which also agrees with previous findings (Melián et al., 2012). Figure 3 shows the mean incipient species size distribution for three different mutation rates. Low mutation rates produce low richness with few highly abundant species, i.e. highly positively skewed abundance distribution. Higher mutation rates (*µ* = 5 × 10*^−^*^3^) tend to generate higher richness and less skewed species abundance distributions. The formation of species always follows the condition *q_min_* > *Q*∗, where *Q*∗ is the mean genetic similarity of the matrix *Q* at equilibrium, as expected from analytical results of Melián et al. (2012). The consideration of plant-animal interactions and morphological constraints for plant reproduction, does not produce qualitatively different results in terms of species abundance distributions compared to previous models (Melián et al., 2012), that did not consider other mating constraints.

**Figure 3:**
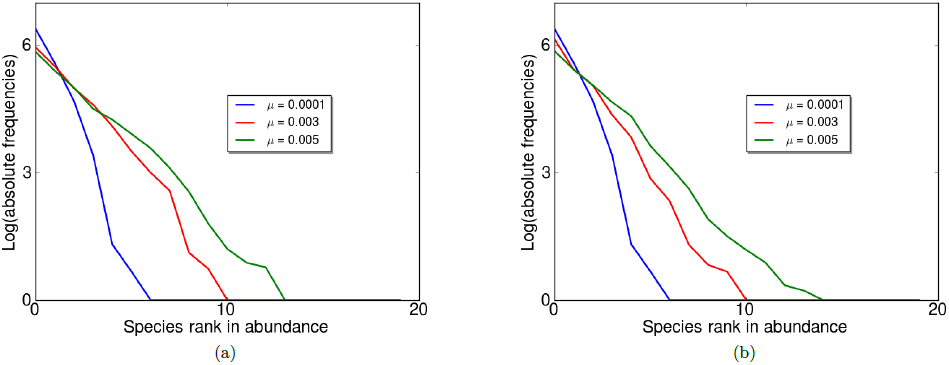
Mean rank abundance distribution of animals (a) and plants (b) after 500 replicates for different mutation rates: *µ* = 5 × 10*^−^*^3^ (green solid line), *µ* = 3 × 10*^−^*^3^(red solid line) and *µ* = 10*^−^*^4^ (blue solid line). Parameters used: *q_min_* = 0.97, *d_max_* = *d^PA^_max_*= 0.3 and *J_P_* = *J_A_* = 10^3^.

### Genotype-phenotype relationship

The genotype-phenotype (G-P) relationship is highly positive as expected from the equation *z* = *g* + *∈*. Figure 4 shows a scatter plot of the G-P relationship of all pairs of individuals which contains three main clouds of points: 1) pairs of individuals of the same species with high genetic (*q_ij_* > *q_min_*) and phenotypic (*p_ij_* > 0.9) similarity, 2) pairs of individuals of the same species with genetic similarity below *q_min_* (*q_ij_* < *q_min_* = 0.97) and high phenotypic similarity (*p_ij_* > 0.9) these are incompatible individuals for mating and yet high phenotypic similarity *p_ij_* > 0.9 and 3) highly genetically dissimilar individuals from different species (*q_ij_* ≪ *q_min_*), but with the presence of highly phenotypically similar individuals (*p_ij_* > 0.9). This is an indication of evolutionary convergence in plants and animals. An increase in mutation rate increases the genetic divergence between species, as expected, but it does not change the G-P relationship qualitatively (see Figure 4). Naturally, we also find pairs of individuals with low genetic (*q_ij_* ≪ *q_min_*) and phenotypic (*p_ij_* < 0.9) similarity.

**Figure 4:**
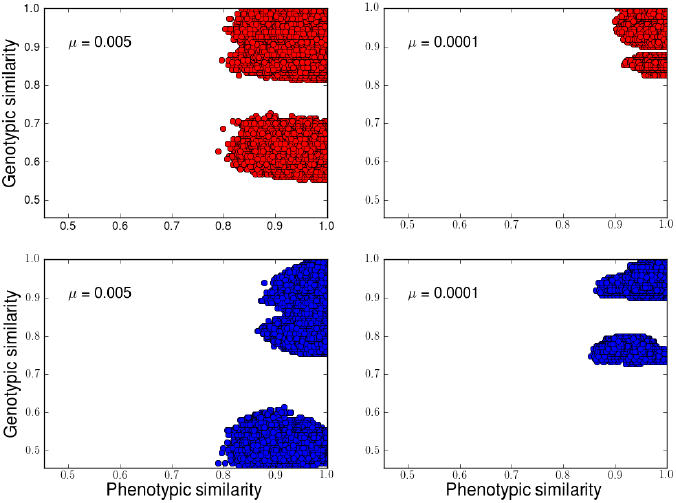
The effect of mutation rate on the genotype-phenotype (G-P) relationship. Top panels show the G-P relationship for animals (red) and bottom panels for plants (blue). Right panels show the G-P relationship for mutation rate *µ* = 5 × 10*^−^*^3^ and left panels for *µ* = 10^−4^. Each plot is an scatter plot, where each filled circle represents phenotypic (*p*_*i*≠*j*_, x-axis) and genetic (*q*_*i*≠*j*_, y-axis) similarity between two individuals of a particular guild (plant or animal) from one replicate. The G-P correlation can be very positive or close to zero depending on the individuals compared. Individuals with high phenotypic similarity and genetic dissimilarity suggests evolutionary convergence of traits, regardless of mutation rate. Parameters used: *q_min_* = 0.97, *d_max_* = *d^PA^_max_* = 0.3 and *J_P_* = *J_A_* = 10^3^.

### Evolutionary convergence and complementarity

The plant and animal trait distributions (i.e. corolla and proboscis lengths) change dramatically during the simulation because of the speciation process, which generates changes in the trait distribution on both guilds resulting in a bimodal distribution (Figure 6). Figure 6 also shows an example of the variation of traits at the species level, where several species of the same guild (plant or animal) can have highly similar trait values (i.e. evolutionary convergence). The figure also shows that there are similar trait values between plant and animal species (i.e. evolutionary complementarity).

**Figure 6:**
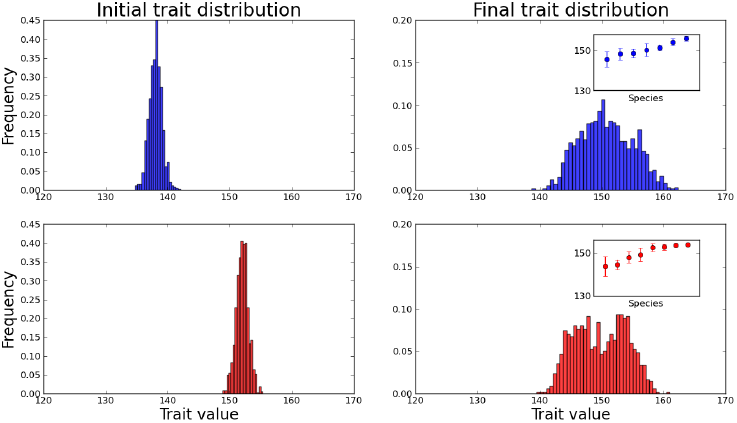
Changes in trait (i.e. phenotype) distribution of plants and animals for a typical replicate simulation. The top panels show the changes in animal (red) trait distribution and bottom panels the changes in plant (blue) trait distribution. Left panels show the initial trait distribution and right panels the final trait distribution. The insets in the right panels show the mean and standard error deviations of traits. The trait distribution changes completely from the initial distribution towards a bimodal distribution in both guilds. Parameters used: *q_min_* = 0.97, *d_max_* = *d^PA^_max_* = 0.3, *µ* = 5 × 10*^−^*^3^ and *J_P_* = *J_A_* = 10^3^.

Evolution of convergence and complementarity occurs in all replicate simulations. Evolutionary convergence appears on average in 17.3 *±* 6% of all possible cases. The number of convergences mainly depends on the number of sister species pairs, i.e. an increase of the number of sister species pairs will decrease the possible number of convergences. Evolutionary complementarity appears with a similar frequency in each replicate simulation, but with a larger variation (20 ± 18%) than convergence. Complementarity mainly depends on the number of plant and animal species; therefore the variation in the number of species at the steady-state between guilds affects the number of complementarity events. An example of evolutionary convergence and complementarity of one replicate is shown in Figure 8.

**Figure 8:**
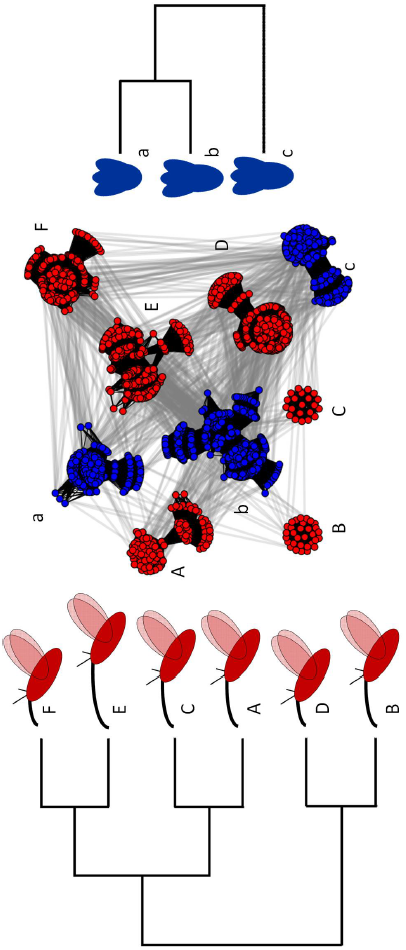
Evolutionary convergence and complementarity in plant-pollinator networks. Cluster trees at the top and the bottom, show genetic similarities between plant (blue) and animal (red) species, respectively. The average species trait, proboscis and corolla length, is sketched with cartoons next to their respective position in the cluster trees. Animals, composed of six species, have two convergent trait events (species A-B, A-C and F-D), while plants, composed of three species, only have one convergent event (species b-c). The central figure shows the network of plant-animal interactions, where each node (colored filled circles) represents an individual in the metacommunity. The network is composed of two types of links: genetic relatedness links (black solid lines) forming clusters that represent species and plant-animal individual-based interaction links (gray lines). The network shows variability in terms of genetic relatedness and plant-animal interactions within a species (i.e. high intraspecific variability). This figure is an example from one replicate simulation. Parameters used: *q_min_* = 0.97, *d_max_* = *d^PA^_max_* = 0.3, *µ* = 5 × 10*^−^*^3^ and *J_P_* = *J_A_* = 10^3^.

### Network topology

The network at the species level is highly nested 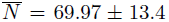 as in real plant-pollinator networks and medium level connectance 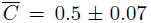 (Figure 5). At the individual level, there is high centrality for a low number of individuals in the whole network regardless of species differences. High centrality also occurs at the intraspecific level. Most individuals in the network have low degree centrality (*DC* ≤ 0.01) and only few individuals have high DC (*DC* ≫ 0.01 ∼ 100 links), which is shown by the positive skewness of the DC distribution (see table 4) and the intraspecific variation (see Figure 7). The betweenness centrality (BC) distribution follows a similar pattern of high intraspecific variation, but less interspecific variation (see Figure 7). The BC distribution is more positively skewed than the DC distribution (see table 4) because of the presence of few individuals with very high BC (*BC* > 0.04), this is more evident in plants than animals. Therefore, the distribution of DC and BC shows that only very few individuals serve as connectors or bridges between species of the opposite guild. High average BC is positively correlated with species abundance (*R*^2^ = 0.798, *p* < 0.001) and partner diversity (i.e. number of partner species) (*R*^2^ = 0.54, *p* < 0.001). Also, there is a high correlation between centrality metrics for both plants and animals (see Table 3). This clearly indicates the importance of these individuals for maintaining the cohesiveness in the network. Although, plants and animals only show slight differences in terms of centrality (see table 4), plant centrality metrics distributions are more asymmetric. The higher asymmetry might be caused by the morphological constraint on plant reproduction. Most individuals have medium levels of closeness centrality 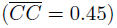. Interestingly, most individuals have neighboring peers in the network that are also interacting with the same mutualistic partners, as indicated by the high level of node redundancy (*NR* > 0.5). However, a few individuals, especially those with high BC, have low node redundancy (*NR* < 0.3) and their extinction could greatly affect network topology. Only the correlation between NR and DC is significantly positive for plants, but not for animals (see Table 3). The average clustering coefficient is higher in plants (*ACC* = 0.053) than animals (*ACC* = 0.038), which suggests that plant network topology tend to be more modular or compartmentalized than animal network topology.

**Figure 5:**
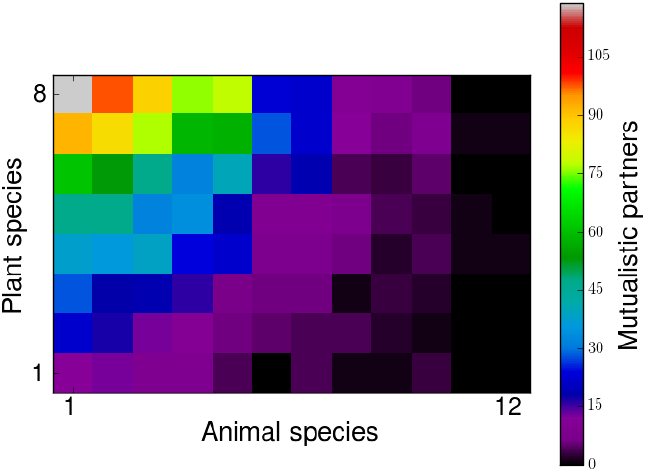
Plant-animal species interaction network. Plant species are represented in rows and animal species in columns. The color gradient indicates the number of mutualistic partners (i.e. individuals interacting) shared between plant and animal species. This matrix comes from one replicate with nine plant and twenty animal species. The network shows high level of nestedness (*N* = 0.72) and intermediate level of connectance (*C* = 0.5). Parameters used: *q_min_* = 0.97, *d_max_* = *d^PA^_max_* = 0.3, *µ* = 5 × 10*^−^*^3^ and *J_P_* = *J_A_* = 10^3^.

**Figure 7:**
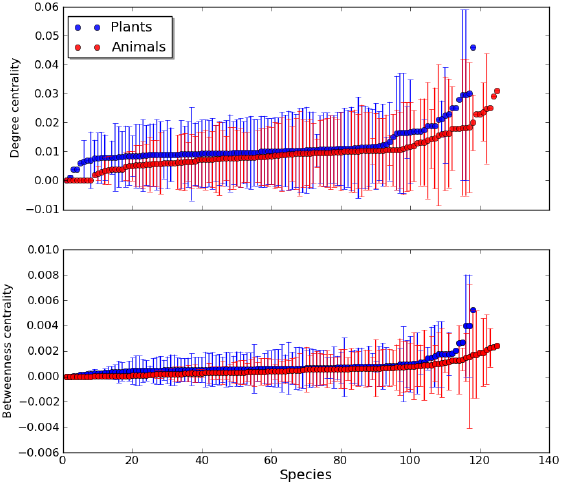
Variation of centrality measurements between and within species. The top panel shows the variation of degree centrality (DC) and the bottom pannel shows the variation of betweenness centrality (BC). Each filled circle represents the average value of DC and BC for one species with their respective standard deviation (vertical thin lines) for plants (blue) and animals (red). The plots represent a sample of 50 replicates. The interspecific and intraspecific variation of BC and DC is quite heterogeneous; a few species tend to have higher BC (> 0.001) and DC (> 0.02) with a large intraspecific variation.

**Table 1:** Glossary of mathematical notation

| Notation | Definition |
| --- | --- |
| $d_{ij}$ | Geographical distance between individual $i$ and $j$ of the same guild |
| $d_{max}$ | Maximum geographical distance to find a mating partner and dispersal |
| $D$ | Geographic distance matrix containing all the $d_{ij}$ values for a guild |
| $d_{ik}^{PA}$ | Geographical distance between plant individual $i$ and animal $k$ |
| $d_{max}^{PA}$ | Maximum geographical distance to find a mutualistic partner |
| $D_{PA}$ | Geographic distance matrix containing all the $d_{ik}^{PA}$ values |
| $q_{ij}$ | Genetic similarity between individual $i$ and $j$ |
| $Q$ | Genetic similarity matrix containing all the pairwise similarity $q_{ij}$ values |
| $q_{min}$ | Minimum genetic similarity above which $i$ and $j$ belong to the same species |
| $p_{ij}$ | Phenotypic similarity between individual $i$ and $j$ |
| $P$ | Phenotypic similarity matrix containing all the $p_{ij}$ values |
| $\mu$ | Mutation rate per locus |
| $N_e$ | Effective population size |
| $z_i$ | Quantitative trait value of individual $i$ |
| $L$ | Size of the genome |
| $\epsilon$ | Environmental effect sampled from a Normal distribution of $\mu_\epsilon = 0$ and $\sigma^2 = 1$ |
| $g_i$ | Genetic effect of an individual $i$ calculated as the sum of alleles across the genome |
| $\hat{p}_{kh}$ | Average phenotypic similarity between species $k$ and $h$ |
| $P_s$ | Phenotypic similarity matrix containing all $\hat{p}_{kl}$ values |
| $P^{PA}$ | Phenotypic similarity matrix between plant and animal species |
| $\hat{q}_{kh}$ | Average genetic similarity between species $k$ and $h$ |
| $Q_s$ | Genetic similarity matrix containing all $\hat{q}_{kl}$ values |

**Table 2:**
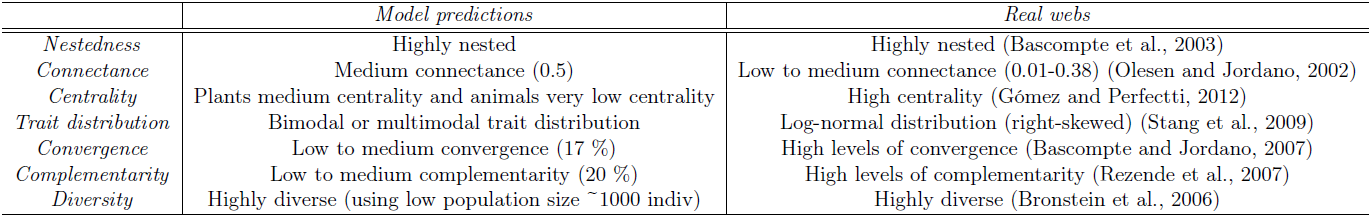
Predictions of the model and observed values in real mutualistic webs. Overall, qualitative predictions are very similar to observed ecological and evolutionary patterns. However, quantitatively we find many differences in the network topology, trait distribution and evolutionary patterns.

**Table 3:**
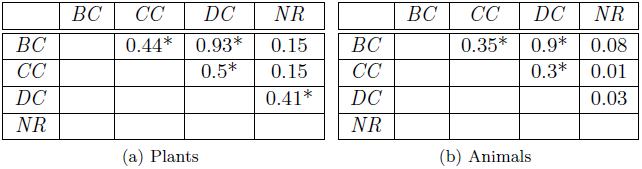
Correlations between centrality metrics and between centrality and node redundancy. BC: Betweenness centrality, CC: closeness centrality, DC: degree centrality, NR: node redundancy. * (p<0.001). Most correlations between centrality metrics are significantly positive. However, correlations between node redundancy and centrality metrics was only significant in plants degree centrality (DC).

|  | *BC* | *CC* | *DC* | *NR* |  | *BC* | *CC* | *DC* | *NR* |
| --- | --- | --- | --- | --- | --- | --- | --- | --- | --- |
| *BC* |  | 0.44* | 0.93* | 0.15 | *BC* |  | 0.35* | 0.9* | 0.08 |
| *CC* |  |  | 0.5* | 0.15 | *CC* |  |  | 0.3* | 0.01 |
| *DC* |  |  |  | 0.41* | *DC* |  |  |  | 0.03 |
| *NR* |  |  |  |  | *NR* |  |  |  |  |

(a) Plants
(b) Animals

**Table 4:** Topology of individual-based plant-pollinator network. Node redundancy (NR) and three centrality metrics were calculated: degree centrality (DC), closeness centrality (CC) and betweenness centrality (BC). The estimates show the skewness, mean and standard error values of the calculated distributions for each metric. The calculations were made for each guild, plants and animals, considering all individuals regardless of species differences. Parameters used: *q_min_* = 0.97, *d_max_* = *d^PA^_max_*= 0.3 and *J_P_* = *J_A_* = 10^3^.

|  | PLANTS |  | ANIMALS |  |
| --- | --- | --- | --- | --- |
|  | MEAN±STD | SKEWNESS | MEAN±STD | SKEWNESS |
| *DC* | 0.01±0.012 | 1.87 | 0.01±0.013 | 2.08 |
| *CC* | 0.45 ± 0.045 | -0.97 | 0.45 ± 0.044 | 0.11 |
| *BC* | 0.00058 ± 0.0013 | 4.47 | 0.00058 ± 0.0014 | 4.78 |
| *NR* | 0.56 ± 0.39 | -0.37 | 0.53 ± 0.4 | -0.38 |

## Discussion

Considering main evolutionary forces in the study of community assembly is crucial to understand the emergence of observed ecological and evolutionary patterns. Several theoretical studies have investigated the evolution of ecological communities assuming niche-related processes as the main drivers of community structure and diversity (Caldarelli et al., 1998; Loeuille and Loreau, 2005; Ingram et al., 2009). Nevertheless, models that only consider neutral processes (e.g. dispersal limitation, ecological drift) are also able to reproduce observed patterns of community diversity and structure (Rosindell et al., 2011). A controversial point of neutral theory was the unrealistic assumption of point-mutation speciation (Hubbell, 2001). This point has been later improved in other models by considering a gradual speciation process phenomenologically (i.e. protracted speciation (Rosindell et al., 2010), see also Etienne and Rosindell (2012)) and mechanistically (i.e. by modelling genetics explicitly (Melián et al., 2012; de Aguiar et al., 2009; Melián et al., 2010)). Genetically and spatially explicit neutral models allow a connection between genetics and community ecology. Our model makes this connection by considering an explicit speciation process and its consequences for the diversity and structure of mutualistic networks. Furthermore, this is the first model, to our knowledge, to study the joint evolution of network structure and quantitative traits in mutualistic networks. Our results show the emergence of some observed topological properties of mutualistic webs and the evolution of trait convergence and complementarity (see Table 2).

Similar to previous neutral genetically explicit eco-evolutionary models (Melián et al., 2012; de Aguiar et al., 2009), two important factors of the speciation process are non-random mating (*q_min_*) and dispersal limitation (*d_max_*). These two factors determine the diversity of communities. Interestingly, Kondrashov and Shpak (1998) found that assortative mating alone in the absence of selection is sufficient to create genetic divergence between individuals and finally the formation of species. Naturally, quantitative changes in the three evolutionary forces considered here (mutation, recombination, genetic drift) are main drivers of the genetic variation in our model and it will ultimately affect diversity patterns. Mutation rate alone can already change the speciation dynamics and species abundance distribution as also shown by Melián et al. (2012). However, dispersal limitation is also a very important driver of the speciation dynamics. It basically determines the gene flow in the metacommunity (Lenormand, 2002) and therefore can help to reinforce the speciation process together with non-random mating (Butlin, 1987). Thus, high assortative mating (high *q_min_*) and high dispersal limitation (low *d_max_*) can maximize the diversity (Melián et al., 2012; de Aguiar et al., 2009).

### Convergence, complementarity and drift

Evolutionary convergence, i.e. the independent evolution of similar features in different evolutionary lineages, of traits is observed in all our replicate simulations and with little variation. Evolutionary convergence has been argued to be a product of multispecific coevolutionary processes (‘diffuse coevolution’)(Janzen, 1980; Jordano et al., 2003; Bascompte and Jordano, 2007; Thompson and Cunningham, 2002) and therefore these patterns are molded by similar selective pressures; as shown by Guimaraes et al. (2011). However, our model shows that evolutionary convergence can occur through the action of the non-selective forces of mutation, recombination and genetic drift. This means that random evolutionary change can cause species to become more similar to each other than their ancestors were, as also shown by Stayton (2008). Stayton (2008) simulated evolution along phylogenies according to a Brownian motion model of trait change and demonstrated that rates of convergence can be quite high when clades are diversifying under only the influence of genetic drift. Furthermore, constraints (e.g. developmental constraints) in the production of variation can also lead to convergence. For example, if the variation produced is limited, then unrelated species are likely to produce the same variation, which may then become fixed in the population by genetic drift (Losos, 2011; Stayton, 2008). This a common feature of biological systems because in the evolution of DNA there are only four possible states for a given nucleotide position, and therefore it is likely that distantly related taxa will independently acquire the same change by chance (Losos, 2011). It can also happen in our model because there are many genotypes that can give the same phenotype.

Developmental constraints are a common explanation for the convergence of traits (Solé et al., 2002; Losos, 2011). However, we still know little about how developmental constraints affect convergence. The tinkering of traits by evolutionary forces largely affect developmental pathways (e.g. gene regulatory networks) (Solé et al., 2002). Thus, developmental pathways are not static, but can diverge through time randomly without substantially affecting phenotype. This is called developmental system drift (DSD) (True and Haag, 2001). We argue that DSD might play an important role in the evolution of morphological traits and it must be considered as another level where drift can be acting (Ohta, 2002), for example, by considering random wiring in gene regulatory networks.

Evolutionary complementarity is also consistently observed in our results but with a larger variation than convergence. Complementarity is argued to be the main result of tight coevolution between mutualistic species by mechanisms, such as trait-matching (e.g. corolla length-proboscis length) (Jordano et al., 2003). There is empirical (Anderson and Johnson, 2008) and theoretical evidence (Gomulkiewicz et al., 2000) for coevolutionary hot spots (Thompson, 1999), which suggests that local selective regimes can promote the coevolution of traits (Ferdy et al., 2002; Bronstein et al., 2006; Gomulkiewicz et al., 2000, 2003; Jordano et al., 2003; Thompson, 2009; Thompson and Cunningham, 2002; Jones et al., 2009). However, we show that low to medium levels of complementarity and convergence can be the product of neutral processes occurring at several levels (i.e. genome, development).

### Evolution of quantitative trait distribution

Our model predicts that the distribution of traits, regardless of species differences, generally evolves towards a bimodal distribution of phenotypes. This result was previously obtained by Kondrashov and Shpak (1998), who assumed absence of selection and assortative mating in a infinite population. Their results show the evolution of traits into two phenotypic classes. Strong assortative mating produces high correlations of allelic effects among all loci, which leads to the evolution of two phenotypic classes: one with alleles increasing the trait and the other with alleles decreasing the trait (Crow and Kimura, 1970). Devaux and Lande (2008) found similar results using a finite diploid population with multiple alleles per locus and they showed that the splitting of the phenotype distribution is possible under strong assortative mating and genetic drift, but the distribution is transient rather than permanent. However, our distribution is not transient, and this is probably because we only considered two allelic states for each locus. As Devaux and Lande (2008) explained, by assuming a normal distribution of allelic effects at each locus we could obtain a more continuous unimodal (i.e. normal) distribution of phenotypes. We need further analytical exploration to thoroughly understand the determinants of trait distributions.

We find a gradient of a species phenotypes from low to high average values (Figure 6). Therefore, a whole spectrum of species phenotypes can emerge in the metacommunity by stochastic processes. However, the predicted trait distribution is not right-skewed as observed in real plant-pollinator communities (Stang et al., 2009) (see Table 2). This might be due to the influence of other forbidden links (e.g. body size) and developmental constraints not considered in this model.

### Neutrality in mutualistic networks: patterns and processes

The morphological constraint for plant reproduction does not seem to change the diversity patterns in plants compared to animals (Figure 3). This suggests that considering this ‘forbidden link’ has no effect on the speciation dynamics of mutualistic networks. However, the topology of plants and animals seems to be slightly different in terms of centrality metrics (i.e. degree and betweenness centrality). Plant topology is more asymmetric than animal topology, which is probably due to the morphological constraint for plant reproduction. This supports the idea that size thresholds of plant and animal mutualistic traits and species abundances promote asymmetry in mutualistic networks (Stang et al., 2009, 2006). The presence of individuals with high centrality (BC and DC) in high-resolution plant-pollinator webs has been found to be related to the fitness of individuals and probably related to specific phenotypes (Gómez and Perfectti, 2012). In our results, high centrality in some plant individuals might be related to the morphological constraint in plant reproduction. Furthermore, higher average clustering coefficient seems higher in plants than animals and it is also probably related to the morphological constraint. This suggests that modularity observed in real mutualistic webs, here indicated by higher average clustering, can be partly an outcome of biological constraints (i.e. forbidden links) (Olesen et al., 2007).

Connectance values are close to the predictions of other neutral network models (Canard et al., 2012) with similar diversity values. However, compared to real mutualistic networks with similar diversity as ours (24 plant and animal species on average), our connectance (*C* = 0.5) is higher than reported webs (*C* = 0.28) (Olesen and Jordano, 2002). This difference in connectance values might also be due to other forbidden links, such as phenology (Olesen et al., 2010).

Nestedness values are also very high, as in real mutualistic networks. The influence of stochastic eco-evolutionary processes and the morphological constraint seems to predict realistic values. However, we think that stochastic processes are more important in determining nestedness. This is based on previous neutral models (Krishna et al., 2008; Canard et al., 2012), which suggests that random interactions, dispersal limitation and species abundance distribution (‘neutral forbidden links’ (Canard et al., 2012)), are determinants of the structure of mutualistic networks.

### Future directions

We have only explored a limited range of the parameter space. For example, we could still explore the effects of genetic similarity (*q_min_*) and spatial structure (*d_max_*, *d^PA^_max_*) on the diversity and structure of mutualistic webs. Based on previous models (Melián et al., 2012; de Aguiar et al., 2009) we expect changes in the diversity of the metacommunity. For example, high values of *q_min_* and shorter geographical distances for mating (*d_max_*) should generate a higher diversity in the metacommunity (Melián et al., 2012). However, extremely low geographic distances for mating could decrease the diversity due to the difficulty of finding mates (Allee effect), especially for high *q_min_* levels.

In our model, assortative mating and the morphological trait are determined for the same multiple loci (i.e. they have the same genetic basis) and these genes show pleiotropic effects. Assortative mating and morphological traits are calculated in a similar way: the sum of genetic differences. This closed relationship between nonrandom mating and an ecological trait is similar to the concept of ‘magic’ traits. A ‘magic’ trait combines a trait subject to divergent selection and another trait related to nonrandom mating (i.e. reproductive isolation) that are pleiotropic expressions of the same gene(s) (Servedio et al., 2011). However, we cannot regard our trait as ‘magic’ because of the absence of disruptive selection forces. There are other alternatives for this relationship between assortative mating and the morphological trait (Servedio et al., 2011). One alternative is that assortative mating and the morphological trait are determined by different sets of genes and express different levels of pleiotropic effects (i.e. a partly ‘magic’ trait (van Doorn and Weissing, 2001)).

One might also explore further the influence of the morphological constraint in the evolution of traits. This constraint might be exerting a weak selection force on the evolution of plant traits in our model. The comparison with other models without any morphological constraint (i.e. only non-random mating) and with morphological constraints for animals and plant reproduction (i.e. phenotypic match), might elucidate the importance of morphological constraints in the evolution of the network.

The possibility to test model predictions with high-resolution data is one of the most important advantages of our model. Plant and pollinator species abundance data, intraspecific trait variation, genetic data, spatial distribution can all be used to test model predictions. Accounting for intraspecific variation helps explaining emergent properties of ecological networks and evolutionary patterns (Bolnick et al., 2011).

We conclude that simple processes (dispersal, demography, mutation, recombination and morphological constraints) can reproduce very well the observed network structure and quantitative trait evolutionary patterns in plant-animal mutualisms.

## Acknowledgements

We thank Martina Stang for discussions.

## Footnotes

^1^defined as reciprocal evolutionary change between species

^2^Parallel evolution is the development of a similar trait in related, but distinct, species descending from the same ancestor

## References

Almeida-Neto, M., P. Guimaraes, P. Guimaraes, R. Loyola, and W. Ulrich. 2008. A consistent metric for nestedness analysis in ecological systems: reconciling concept and measurement. Oikos 117:1227–1239.

Anderson, B., and S. Johnson. 2008. The Geographical mosaic of coevolution in a plant-pollinator mutualism. Evolution 62:220–225.

Bascompte, J., and P. Jordano. 2007. Plant-animal mutualistic networks: The architecture of biodiversity. Annu. Rev. Ecol. Evol. Syst. 38:567–593.

Bascompte, J., P. Jordano, C. J. Melian, and J. M. Olesen. 2003. The nested assembly of plant-animal mutualistic networks. Proc. Natl. Acad. Sci. USA 100:9383–9387.

Bastolla, U., M. Fortuna, A. Pascual-García, A. Ferrera, B. Luque, and J. Bascompte. 2009. The architecture of mutualistic networks minimizes competition and increases biodiversity. Nature 458:1018–1020. URL http://dx.doi.org/10.1038/nature07950.

Bolnick, D., P. Amarasekare, M. Araújo, R. Bürger, J. Levine, M. Novak, V. Rudolf, S. Schreiber, M. Urban, and D. Vasseur. 2011. Why intraspecific trait variation matters in community ecology. Trends Ecol Evol 26:183–192. URL http://www.sciencedirect.com/science/article/pii/S0169534711000243.

Borgatti, S. 2005. Centrality and network flow. Social Networks 27:55–71.

Borgatti, S., and D. Halgin, In press. The Sage Handbook of Social Network Analysis, Chapter analyzing affiliation networks. Sage Publications.

Bronstein, J. L., R. Alarcon, and M. Geber. 2006. The evolution of plant-insect mutualisms. New Phytol. 172:412–428.

Butlin, R. 1987. Speciation by reinforcement. Trends Ecol Evol 2:8–13.

Caldarelli, G., P. Higgs, and A. McKane. 1998. Modelling coevolution in multispecies communities. Journal Theoretical Biology 193:345–358.

Canard, E., N. Mouquet, L. Marescot, K. Gaston, D. Gravel, and D. Mouillot. 2012. Emergence of structural patterns in neutral trophic networks. Plos One 7:e38295.

Crow, J., and M. Kimura. 1970. An introduction to population genetics theory. Harper & Row Publishers, New York.

Darwin, C. 1862a. Fertilisation of Orchids. Murray.

Darwin, C. 1862b. On the Various Contrivances by Which British and Foreign Orchids are Fertilised by Insects, and on the Good Effects of Intercrossing. Murray.

de Aguiar, M., M. Baranger, E. Baptestini, L. Kaufman, and Y. Bar-Yam. 2009. Global patterns of speciation and diversity. Nature 460:384–387.

Devaux, C., and R. Lande. 2008. Incipient allochronic speciation due to non-selective assortative mating by flowering time, mutation and genetic drift. Proc R Soc B 275:2723–2732.

Etienne, R., and J. Rosindell. 2012. Prolonging the past counteracts the pull of the present: protracted speciation can explain observed slowdowns in diversification. Systematic Biology 61:204–213.

Ferdy, J., L. Despres, and B. Godelle. 2002. Evolution of mutualism between globeflowers and their pollinating flies. Journal of Theoretical Biology 217:219–234.

Ferriere, R., M. Gauduchon, and J. L. Bronstein. 2007. Evolution and persistence of obligate mutualists and exploiters: competition for partners and evolutionary immunization. Ecology Letters 10:115–126.

Goh, K.-I., E. Oh, B. Kahng, and D. Kim. 2003. Betweenness centrality correlation in social networks. Phys. Rev. E 67:017101.

Gómez, J., F. Perfectti, and P. Jordano. 2011. The Functional consequences of mutualistic network architecture. PloS ONE 6:e16143.

Gómez, J. M., and F. Perfectti. 2012. Fitness consequences of centrality in mutualistic individual-based networks. Proc R Soc B 279:1754–1760.

Gomulkiewicz, R., S. L. Nuismer, and J. N. Thompson. 2003. Coevolution in variable mutualisms. The American Naturalist 162:S80–S93.

Gomulkiewicz, R., J. N. Thompson, R. D. Holt, S. L. Nuismer, and M. E. Hochberg. 2000. Hot spots, cold spots, and the geographic mosaic theory of coevolution. The American Naturalist 156:156–174.

González, A. M., B. Dalsgaard, and J. Olesen. 2010. Centrality measures and the importance of generalist species in pollination networks. Ecological Complexity 7:36–43.

Guimaraes, P., P. Jordano, and J. Thompson. 2011. Evolution and coevolution in mutualistic networks. Ecol Lett 14:877–885.

Hagberg, A. A., D. A. Schult, and P. J. Swart, 2008. Exploring network structure, dynamics, and function using NetworkX. Pages 11–15 *in* T. V. Gäel Varoquaux and J. Millman, editors. Proceedings of the 7th Python in Science Conference (SciPy2008). Pasadena, CA USA.

Howe, H., and J. Smallwood. 1982. Ecology of seed dispersal. Ann Rev. Ecol. Syst. 13:201–228.

Hubbell, S. 2001. The unified neutral theory of biodiversity and biogeography. Princeton University Press, Princeton, NJ.

Huerta-Cepas, J., J. Dopazo, and T. Gabaldón. 2010. ETE: a python environment for tree exploration. BMC Bioinformatics 11:24.

Hunter, J. 2007. Matplotlib: A 2D graphics environment. Computing In Science & Engineering 9:90–95.

Ingram, T., L. Harmon, and J. Shurin. 2009. Niche evolution, trophic structure, and species turnover in model food webs. The American Naturalist 174:56–57.

Janzen, D. 1980. When is it coevolution? Evolution 34:611–612.

Jones, E., R. Ferrière, and J. Bronstein. 2009. Eco-evolutionary dynamics of mutualists and exploiters. Am Nat 174:780–794.

Jordán, F., Z. Benedekc, and J. Podanic. 2007. Quantifying positional importance in food webs: A comparison of centrality indices. Ecological Modelling 205:270–275.

Jordano, P., J. Bascompte, and J. Olesen. 2003. Invariant properties in coevolutionary networks of plant-animal interactions. Ecol. Lett. 6:69–81.

Jousselin, E., J.-Y. Rasplus, and F. Kjellberg. 2003. Convergence and coevolution in a mutualism: evidence from a molecular phylogeny of ficus. Evolution 57:1255–1269.

Kimura, M. 1983. The neutral theory of molecular evolution. Cambridge University Press, Cambridge, UK.

Klovdahl, A. 1985. Social networks and the spread of infectious diseases: The AIDS example. Social Science & Medicine 21:1203–1216.

Kondrashov, A., and M. Shpak. 1998. On the origin of species by means of assortative mating. Proc. R. Soc. Lond. B 265:2273–2278.

Krishna, A., P. Guimaraes, P. Jordano, and J. Bascompte. 2008. A neutral-niche theory of nestedness in mutualistic networks. Oikos 117:1609–1618.

Latapy, M., C. Magnien, and N. D. Vecchio. 2008. Basic notions for the analysis of large two-mode networks. Social Networks 30:31–48.

Law, R., J. L. Bronstein, and R. Ferriere. 2001. On mutualists and exploiters: plant-insect coevolution in pollinating seed-parasite systems. Journal of Theoretical Biology 212:373–389. URL http://dx.doi.org/10.1006/jtbi.2001.2383.

Lenormand, T. 2002. Gene flow and the limits to natural selection. Trends Ecol Evol 17:183–189.

Loeuille, N., and M. Loreau. 2005. Evolutionary emergence of size-structured food webs. Proceedings of the National Academy of Sciences of the United States of America 102:5761–5766. URL http://dx.doi.org/10.1073/pnas.0408424102.

Losos, J. 2011. Convergence, adaptation, and constraint. Evolution 65:1827–1840.

Lynch, M. 2007. The frailty of adaptive hypotheses for the origins of organismal complexity. Proc Natl Acad Sci USA 104:8597–8604.

Melián, C., D. Alonso, D. Vázquez, J. Regetz, and S. Allesina. 2010. Frequency-dependent selection predicts patterns of radiations and biodiversity. PLoS Comput Biol 6. URL http://dx.doi.org/10.1371/journal.pcbi.1000892.

Melián, C. A. D, S. Allesina, R. Condit, and R. Etienne. 2012. Does sex speed up evolutionary rate and increase biodiversity? PloS Comput. Biol. 8:e1002414.

Newman, M. 2003. The structure and function of complex networks. SIAM Rev. 45:167.

Ohta, T. 2002. Near-neutrality in evolution of genes and gene regulation. Proc Natl Acad Sci USA 99:16134–16137.

Okuyama, T., and J. N. Holland. 2008. Network structural properties mediate the stability of mutualistic communities. Ecol. Lett. 11:208–216.

Olesen, J., J. Bascompte, Y. Dupont, H. Elberling, C. Rasmussen, and P. Jordano. 2010. Missing and forbidden links in mutualistic networks. Proc. R. Soc. Lond. B URL http://dx.doi.org/10.1098/rspb.2010.1371.

Olesen, J., and P. Jordano. 2002. Geographic patterns in plant-pollinator mutualistic networks. Ecology 89:2416–2424.

Olesen, J. M., J. Bascompte, Y. L. Dupont, and P. Jordano. 2007. The modularity of pollination networks. Proc. Natl. Acad. Sci. USA 104:19891–19896.

Pérez, F., and B. Granger. 2007. IPython: a System for Interactive Scientific Computing. Comput. Sci. Eng. 9:21–29.

Rezende, E. L., P. Jordano, and J. Bascompte. 2007. Effects of phenotypic complementarity and phylogeny on the nested structure of mutualistic networks. Oikos 116:1919–1929.

Rosindell, J., S. J. Cornell, S. P. Hubbell, and R. S. Etienne. 2010. Protracted speciation revitalizes the neutral theory of biodiversity. Ecol Lett 13:716–727.

Rosindell, J., S. Hubbell, and R. Etienne. 2011. The unified neutral theory of biodiversity and biogeography at age ten. Trends Ecol Evol 26:340–348.

Servedio, M., G. V. M. Kopp, A. Frame, and P. Nosil. 2011. Magic traits in speciation: ‘magic’ but not rare? Trends Ecol Evol 26:389–399.

Solé, R., R. Ferrer-Cancho, J. Montoya, and S. Valverde. 2002. Selection, tinkering, and emergence in complex networks. Complexity 8:20–33.

Stang, M., P. Klinkhamer, and E. van der Meijden. 2006. Size constraints and flower abundance determine the number of interactions in a plant-flower visitor web. Oikos 112:111–121.

Stang, M., P. Klinkhamer, N. Waser, I. Stang, and E. van der Meijden. 2009. Size-specific interaction patterns and size matching in a plant-pollinator interaction web. Ann Bot 103:1459–1469.

Stayton, C. 2008. Is convergence surprising? An examination of the frequency of convergence in simulated datasets. J. Theo. Biol. 252:1–14.

Thompson, J. 2009. The coevolving web of life. The American Naturalist 173:125–140.

Thompson, J. N. 1999. Specific hypotheses on the geographic mosaic of coevolution. American Naturalist 153:S1–S14.

Thompson, J. N., and B. M. Cunningham. 2002. Geographic structure and dynamics of coevolutionary selection. Nature 417:735–738.

True, J. R., and E. S. Haag. 2001. Developmental system drift and flexibility in evolutionary trajectories. Evolution & Development 3:109–119.

van Doorn, G., and F. Weissing. 2001. Ecological versus sexual models of sympatric speciation: a synthesis. Selection 2:17–40.

Vellend, M. 2010. Conceptual synthesis in community ecology. The Quarterly Review of Biology 85:183–206.

Waser, N., L. Chittka, M. Price, N. Williams, and J. Ollerton. 1996. Generalization in pollination systems, and why it matters. Ecology 77:1043–1060.

